# Evolution of intracellular free radical load in colon adenocarcinoma cells over the course of butyrate-induced redifferentiation

**DOI:** 10.1101/2023.03.22.533138

**Authors:** Alina Sigaeva, Eline Zijlema, Yue Zhang, Romana Schirhagl

**Affiliations:** Bioimaging and Bioanalysis Group, Department of Biomedical Engineering, University Medical Center Groningen, Antonius Deusinglaan 1, 9713AV Groningen, the Netherlands

**Keywords:** diamond relaxometry, fluorescent nanodiamonds, free radicals, colon cancer

## Abstract

Fluorescent nanodiamonds have exceptional optical properties and are highly biocompatible, which allows to use them as labels for long-term tracking of the cells. The research fields that make use of this application of nanodiamonds include stem cell biology and cancer biology, where quiescent and differentiating cells can be traced *in vitro* and *in vivo*. However, these studies focus on using nanodiamonds as simple labels, whereas they can serve as highly sensitive intracellular sensors for free radical species. In this study, we aimed to bring the two approaches together and to assess the free radical production in the cells over the course of their differentiation.

We report on the successful enterocytic differentiation of HT-29 colon adenocarcinoma cells, pre-loaded with fluorescent nanodiamonds. The cells were cultured in butyrate-free or butyrate-supplemented medium for 13 days. Butyrate-treated cells developed the morphological and molecular traits, characteristic for normal enterocytes. Fluorescent nanodiamonds did not have a negative effect on the process of differentiation. Moreover, the particles could be found in the cytoplasm of both undifferentiated and re-differentiated cells even after 13 days of culture. The internalized nanodiamonds were used to assess the free radical load in the undifferentiated and re-differentiated HT-29 cells at different stages of the experiment. Consistently with previous findings, re-differentiated HT-29 cells showed higher free radical load than undifferentiated ones.

## 1. Introduction

Over the last decades, fluorescent nanodiamonds (FNDs) – diamond particles with a size range of several nanometers to a hundred nanometers – have been attracting interest of researchers due to their exceptional optical properties. The fluorescence of FNDs stems from their color centers – defects of the crystal structure. The most commonly used defect is nitrogen vacancy (NV) center. As the NV-center fluorescence is not prone to bleaching or blinking, FNDs with this type of defects make a very good substitute for fluorescent dyes, especially if long-term labelling or imaging is required. At the same time, this extremely stable fluorescence can be modulated by external physical factors. The combination of these properties has led to the development of nanodiamond-based quantum sensing protocols, where the fluorescence of FNDs in the sample is monitored to obtain information on parameters as temperature^1,2^, pH^3^, or magnetic fields^4^. One of the most exciting research avenues is applying these sensing protocols to biological samples.

Multiple studies have shown that the nanodiamonds have exceptional biocompatibility, both *in vitro* and *in vivo*^5^. In a proof-of-concept work by Fang et al., FNDs were retained in the cytoplasm of cultured HeLa cells, 489-2.1 multipotent stromal cells, and 3T3-L1 preadipocytes for at least 8 days without causing substantial changes in cell growth and proliferation^6^. In a remarkable work by Mohan et al., FNDs injected into the gonads of *Caenorhabditis elegans* were transported into the worms’ oocytes, which then were successfully fertilized and could develop into normal embryos and larvae^7^.

Stable fluorescence and high biocompatibility of FNDs make these nanoparticles suitable for tracking the cells in complex systems, such as live organisms. FNDs have been used to identify and track quiescent cancer stem cells for 20 days^8^. These cells have very low division rates, which means that internalized FNDs are retained in the same cell and not divided between two daughter cells, resulting in high fluorescence signal coming from quiescent cells. In another study, FND-labelled human mesenchymal stromal cells were injected in miniature pigs. The particles’ fluorescence allowed the researchers to quantitatively detect the transplanted cells in the tissues of the animals 48 hours after the injection^9^.

Liu et al. have shown that FNDs are retained in A459 lung cancer cells and 3T3-L1 cells for at least 10 days without affecting the cell cycle progression and cell division. Moreover, internalized FNDs did not interfere with adipogenic differentiation of 3T3-L1 cells^10^.

Similarly, human mesenchymal stromal cells, isolated from lipoaspirate, could be differentiated into adipocytes after being labelled with FNDs^11^. Wu et al. demonstrated differentiation of FND-loaded human adipose-derived stem/stromal cells into chondrocytes in the xenografts^12^. Hsu et al. have successfully achieved labelling of embryonal carcinoma stem cells and their differentiation into neurons^13^. In an *in vivo* differentiation study, FND-loaded transplanted lung progenitor cells could be identified in murine tissues up to 7 days after transplantation, and the presence of FNDs did not affect the differentiation of progenitor cells into pneumocytes^14^.

Apart from cell labelling and tracking, nanodiamonds have been used as theranostic platforms, particularly in cancer therapy^15^. Conjugates of nanodiamonds and therapeutic agents (e.g., anti-cancer drugs, such as doxorubicin) have higher targeting and retention rates than non-conjugated drugs. This leads to higher efficiency of the therapy, less pronounced side effects, suppression of chemoresistance, and better elimination of cancer stem cells^15^. At the same time, nanodiamonds can facilitate imaging of the tumours in different modalities, such as MRI^16^, fluorescent and photoacoustic imaging^17^. Together, these properties make nanodiamonds a promising and multifunctional tool for cancer diagnostics and therapy. However, the potential of FNDs for revealing the properties of cancer cells and differentiating cells has not been fully realized yet.

In this study, we focus on the application of nanodiamonds in the HT-29 cell line. The HT-29 cell line is a human colon adenocarcinoma cell line. It is commonly used as an *in vitro* model for colon cancer progression, the underlying molecular mechanisms, and response to treatment. Colorectal cancer is one of the most common causes of cancer-related deaths worldwide, as it is often diagnosed at an advanced stage, when the tumours are likely to develop therapy resistance^18^. It is, therefore, important to develop the tools for early diagnostic of colorectal cancer, for assessing the sensitivity of the tumours to the therapy, and for better understanding of the emerging resistance.

HT-29 cells are remarkable, as they can be forced into a more differentiated state, resembling normal enterocytes, if cultured in butyrate-enriched medium^19^. This differentiated state is achieved at 10-12 days of incubation with butyrate. Re-differentiated cells become flattened and more adherent, form junctional complexes, exhibit morphological polarity with distinct apical and basolateral surfaces, and become capable of transepithelial transport and mucus secretion^19^. Butyrate also inhibits the monolayer formation and reduces clonogenicity by >99%. On the ultrastructural level, butyrate-treated cells show a well-defined brush border, characteristic for differentiated epithelial cells^20^. In the gastrointestinal tract, butyrate is produced by resident microorganisms and has been identified as an antineoplastic and antiinflammatory agent^21^.

It has been previously shown that butyrate treatment can induce changes in the redox parameters of HT-29 cells, such as redox potential, free radical levels, production of reactive oxygen species (ROS), reactive nitrogen oxide species (RNOS), and release of hydrogen peroxide^22^. Acute exposure of undifferentiated HT-29 cells to butyrate results in high levels of oxidative stress and cell death, whereas differentiated HT-29 cells release hydrogen peroxide, but do not exhibit signs of stress. Similarly, butyrate triggers ROS production and apoptosis in SW480 cells, another colorectal cancer cell line, but not in normal colon epithelial cells, and can inhibit *in vivo* tumour growth^23^. Interestingly, the baseline free radical load, as measured by electron spin resonance, was higher in differentiated HT-29 cells, as opposed to undifferentiated cells. Together, these findings indicate the potential functional involvement of free radical/ROS production in colonocyte differentiation and colon cancer development. However, it is unknown, how and when this characteristic response is established during the differentiation process. In the current study, we use nanodiamond-based quantum sensing to assess the differences in free radical load between the differentiated and non-differentiated state of the HT-29 cells.

## 2. Materials and methods

### 2.1. Cell line

Human colon adenocarcinoma HT-29 cells were routinely cultured in Dulbecco’s Modified Eagle’s Medium (Cellutron Life Technologies, USA) with a high glucose concentration, 10% fetal bovine serum (FBS, ScienCell, USA), 100 U/mL penicillin, 100 mg/mL streptomycin (Life Technologies) (DMEM-HG medium). The cells were cultured at +37°C, 5% CO2 and were passaged when they reached 70-80% confluency.

### 2.2. Uptake of fluorescent nanodiamonds and butyrate-induced differentiation

Previous research has shown that the uptake of FNDs was increased when the cells first were treated with trypsin-EDTA^24^. In this study, we exposed HT-29 cells to FNDs during passaging. We used 120-nm FNDs (Adamas Nanotechnologies) at the final concentration of 3 μg/mL. To prepare the FND-containing medium, the stock solution (1 mg/mL) of FNDs was diluted in pure FBS. The FBS-FND mixture was then diluted in serum-free DMEM-HG medium, so that the final solution contained 10% of FBS and 3 μg/mL of FNDs.

When the cells have reached a confluency of 70-80%, they were treated with 0.05% trypsin-EDTA (Gibco), until the cells were detached from the bottom of the flask. The cell suspension was then collected and centrifuged at x 1000 rpm for 3 minutes to remove trypsin-EDTA. The supernatant was then discarded, and the cell pellet was resuspended in complete DMEM-HG medium. A small volume of cell suspension was taken for cell counting. Then, the cells were added to either FND-containing medium or FND-free DMEM-HG complete and seeded in quartered glass bottom Petri dishes at 1 x 10^4^ cells/cm^2^ (500 μL cell suspension per quarter). After 48 hours, in the butyrate-treated group, the medium was changed to a medium containing 5 mM sodium butyrate (Sigma-Aldrich). The cells were grown in four different conditions: no FNDs and no sodium butyrate, with FNDs and no sodium butyrate, no FNDs and with sodium butyrate, and with FNDs and sodium butyrate. The medium was changed every 2 days. The cells were cultured for 13 days and collected for staining and T1 measurements at 3, 5, 8, 10 and 13 days.

### 2.3. Cell fixation, immunocytochemistry and imaging

To assess the differentiation status of the cells, we performed immunocytochemical staining for villin – a protein expressed in the brush border of differentiated epithelial cells. For the staining, the cells were rinsed with PBS, then fixed with 3.7% paraformaldehyde (PFA) for 12 minutes, and washed three times with PBS. Next, the samples were incubated with 0.5% Triton X-100 for 10 minutes, followed by washing with PBS 3 times for 5-minute. The sites of non-specific binding were then blocked for 30 minutes in 1% BSA in PBS (PBSA), after which the samples were incubated with the primary anti-villin antibody (1:200 in PBSA, Abcam, ab201989) over night at room temperature. The next day, the samples were washed with PBSA three times for 5 minutes per wash. The cells were incubated with the secondary antibody (goat-anti-mouse, FITC conjugated, 1:200 in PBSA, Jackson ImmunoResearch Europe Ltd, #115-095-146) for 1 hour at room temperature in the dark, followed by washing 2 times with PBSA for 5 minutes. Next, the samples were washed once with PBS for 5 minutes and stored at +4°C in 1% PFA in PBS.

To assess the results of immunostaining, we recorded three images per sample from different areas of the dish. Cell morphology was assessed from the bright field images. Villin expression was estimated from the average fluorescence intensity in the region of interest (ROI - the cell clusters) in the FITC channel. FNDs were identified by far-red fluorescence (excitation wavelength 561 nm, emission range: 650-758 nm).

### 2.4. Nanodiamond relaxometry (T1 measurements)

To assess the free radical production in HT29 cells at different stages of differentiation, we performed diamond relaxometry, making use of the fluorescent defects in internalized nanodiamonds (NV-centers). With this technique, the magnetic noise of the immediate environment of the particle is read out optically. First, the NV-centers are pumped by a green laser pulse into a bright state. Then the fluorescence of the NV-centers is recorded after increasing dark times. During the dark times, the system goes through natural relaxation to the darker equilibrium between the ms=0 and ms=+-1 states, and the NV-center fluorescent signal becomes dimmer. Higher concentrations of free radicals result in higher levels of magnetic noise, which leads to faster relaxation of NV-centers. It means that the fluorescent signal will become dimmer after a shorter dark time, as compared to the system with little magnetic noise (lower concentrations of free radicals). The relaxation time can be described by the T1 constant, which is extracted from the relationship between NV-center fluorescence and the dark time, during which the system was allowed to relax.

The relaxometry measurements were done with a home-made setup, based on a confocal microscope^25,26^. The dark times vary from 200 ns to 1 ms, and the sequence of laser pulses separated by dark times was repeated 1000 times for each T1 measurement. Each measurement lasted approximately 1 minute. To make sure that the particle is not lost during the measurement, the setup performed automated recentering every 3 seconds.

For each stage of differentiation, we obtained 50 T1 measurements, each from a different particle in a different cell within the same sample.

### 2.5. Statistical analysis

The statistical analysis is performed using the Graphpad Prism 8 software. Depending on the data, either a t-test or two-way ANOVA was performed with an alpha value of 0.05. The following P-value indication is used: *P<0.05, **P<0.01 and ***P<0.001.

## 3. Results

### 3.1. Butyrate inhibits cell proliferation and induces changes in the cell morphology

In line with the previous findings, the proliferation rate of HT-29 cells exposed to sodium butyrate was lower than that of non-treated cells. HT-29 cells cultured in butyrate-free medium were dividing throughout the experiment, occupying most of the available dish surface and forming large clusters or monolayers **(Figure 1).** In contrast, butyrate-treated HT-29 cells barely increased in numbers. Even by day 13, we could clearly see small clusters of cells, and no monolayer formation was observed. Moreover, the cell morphology was clearly altered. Butyrate-treated cells became more flattened and spread over the surface, as compared to day 1 of the treatment. These morphological changes are consistent with previously described effects of sodium butyrate on HT-29 cells^19^.

**Figure 1.**
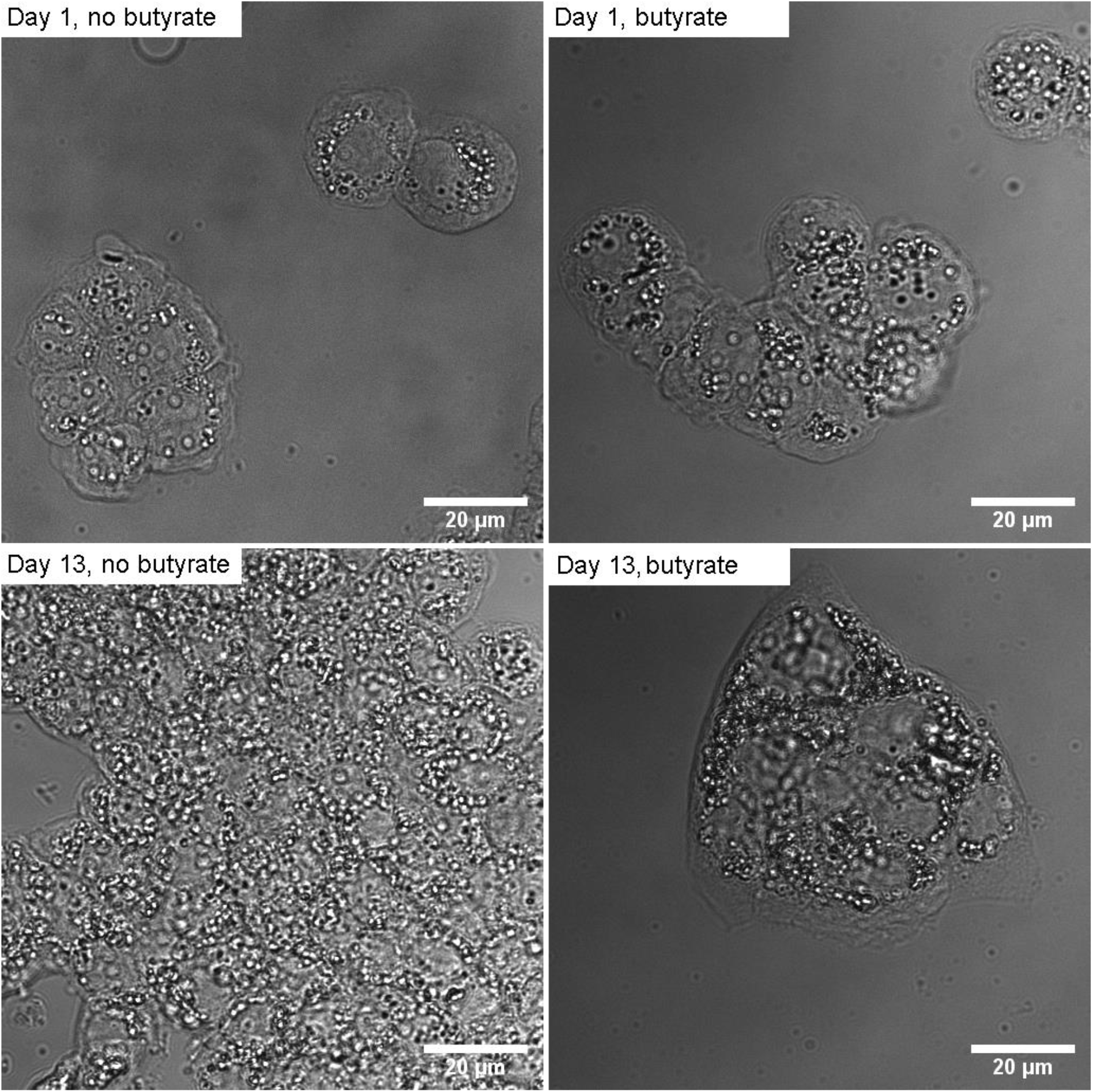
Bright-field (DIC) images of HT-29 cells at different stages of the experiment.

### 3.2. Butyrate-treated cells show higher levels of villin expression

Anti-villin staining confirms that butyrate-treated HT-29 cells become progressively more and more differentiated over the course of the experiment **(Figure 2–6).** In HT-29 cells that did not have FNDs (**Figure 6,** black lines), villin expression was significantly higher in butyrate-treated cells, starting from day 8. At day 13, the p-value of the multiple comparisons drops below statistical significance (p = 0.2613), which might be caused by a small sample size. Similarly, FND-loaded HT-29 cells were showing higher levels of villin expression, when treated with butyrate. In this case, multiple comparisons yielded statistically significant differences only at day 3, possible due to insufficient sample size. In both FND-carrying and FND-free cells, butyrate treatment had very significant effects on the villin expression, as shown by two-way ANOVA **(Table 1).**

**Figure 2.**
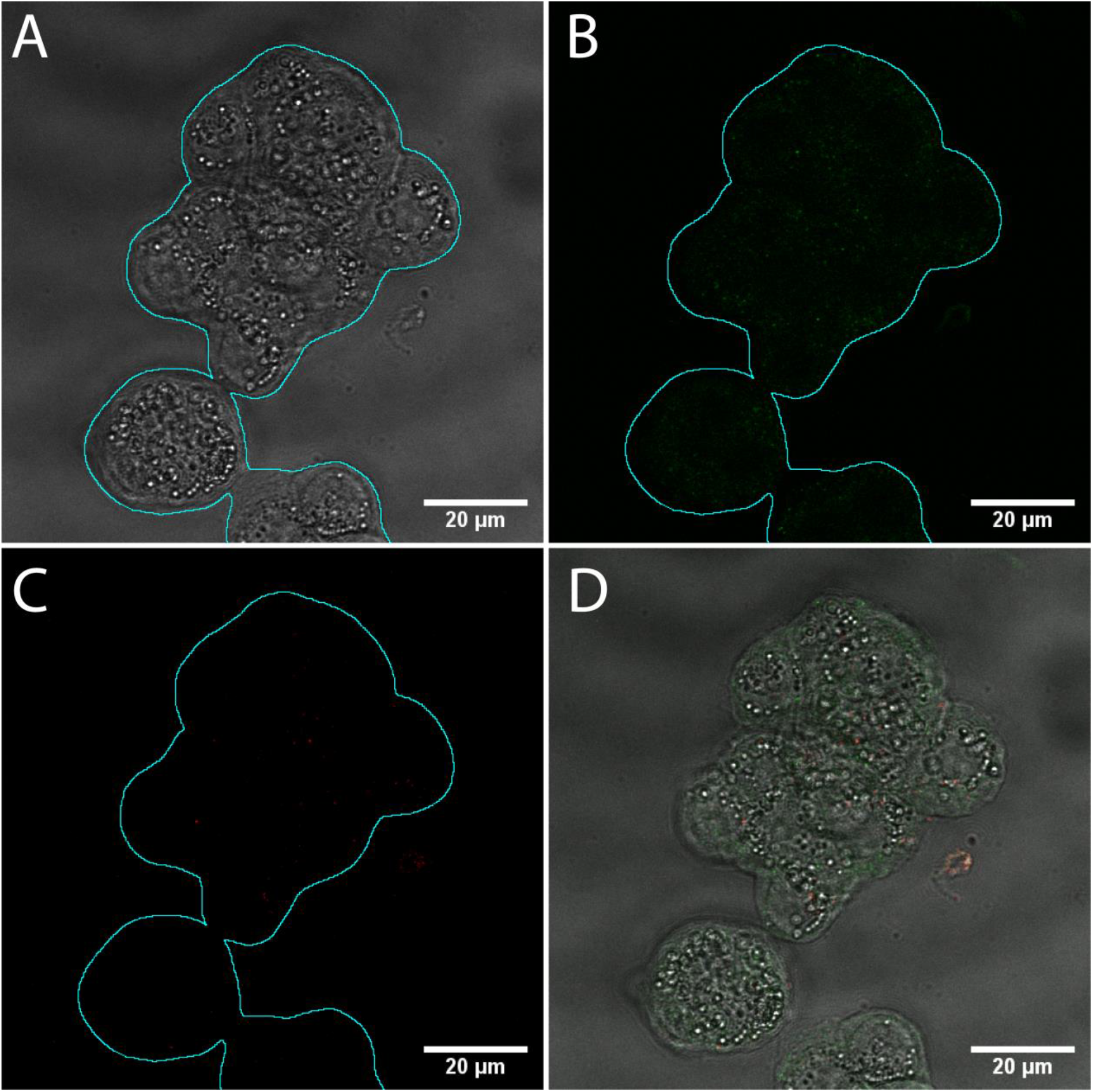
FND-loaded cells, cultured in butyrate-free DMEM-HG, day 1. The DIC image of the cell cluster (A) was used to define the region of interest for future image analysis (cyan outline). Minimal signal was observed in the FITC channel, corresponding to anti-villin staining (B). Internalized FNDs could be clearly seen in the cytoplasm in the far red channel (C). (D) shows the overlay of three channels.

**Figure 3.**
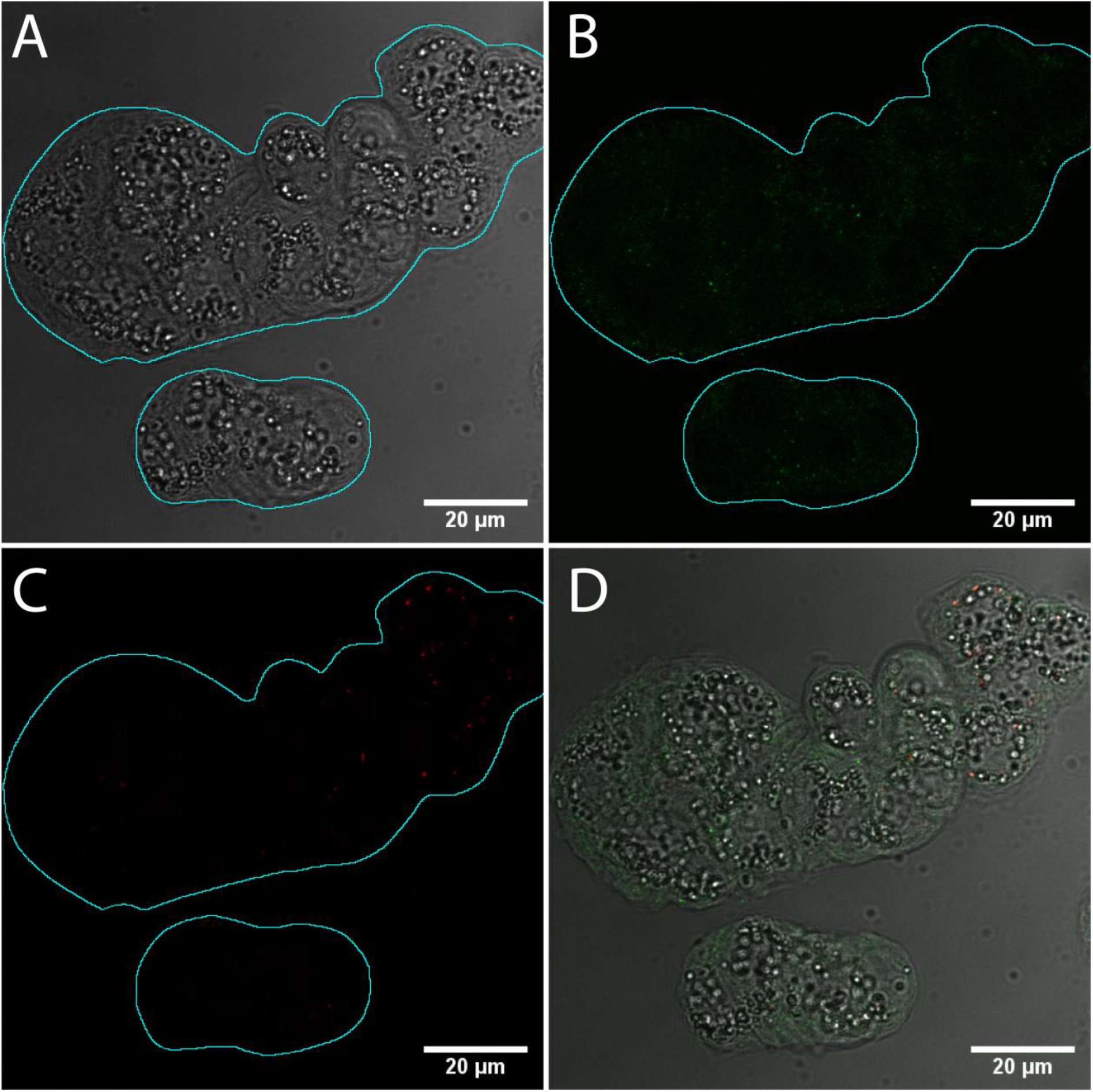
FND-loaded cells, cultured in butyrate-supplemented DMEM-HG, day 1. The DIC image of the cell cluster (A) was used to define the region of interest for future image analysis (cyan outline). Like in **Figure 2,** the villin staining was not pronounced (B), and the cells were carrying FNDs in the cytoplasm (C). (D) shows the overlay of three channels.

**Figure 4.**
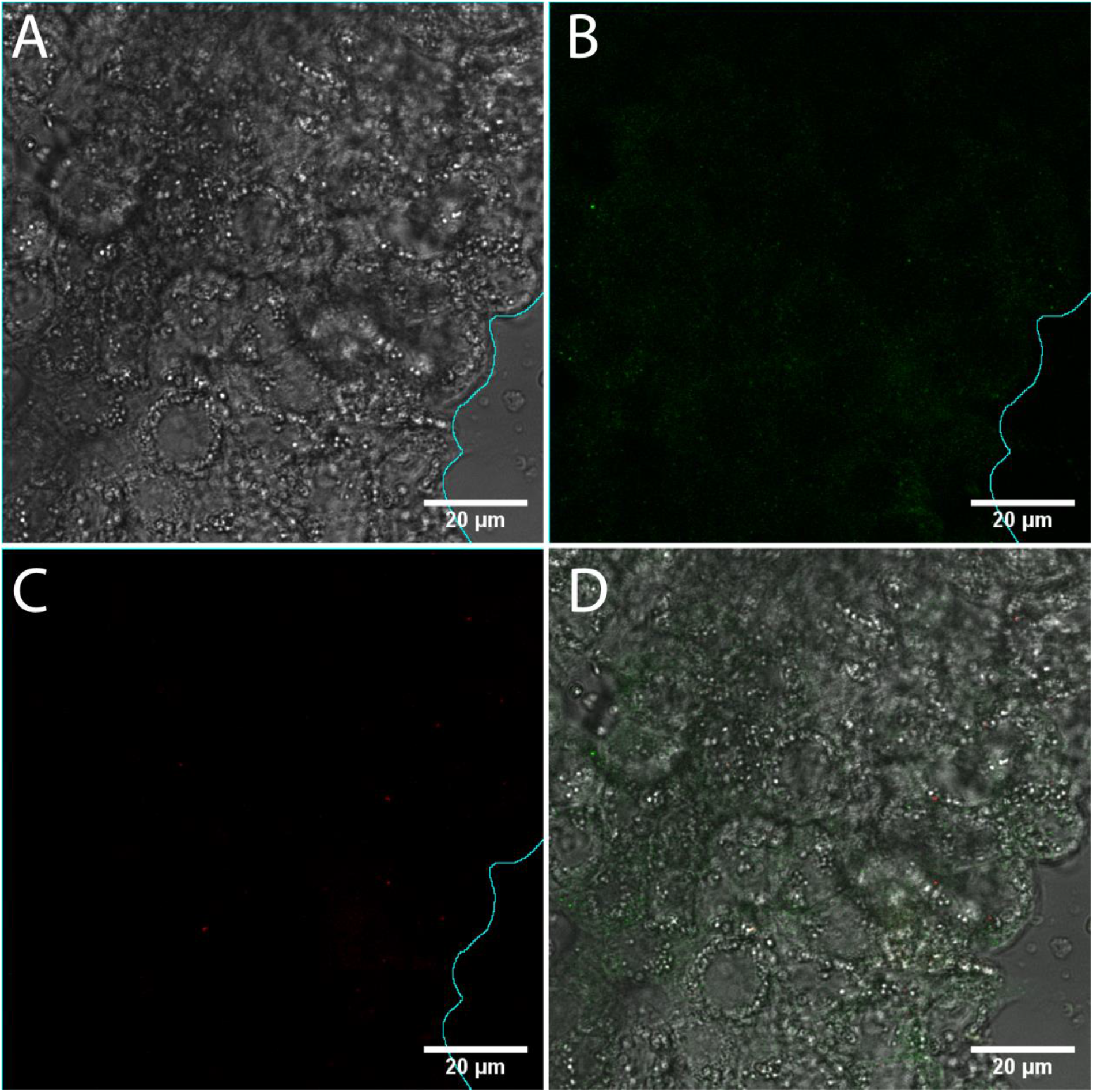
FND-loaded cells, cultured in butyrate-free DMEM-HG, day 13. The DIC image (A) shows a large cluster of cells, close to a monolayer, with some cells growing on top of the others. In the FITC channel (B), there is still little signal within the region of interest (cyan outline). FNDs can be detected in the cytoplasm of the cells (C), but their counts are clearly lower than in **Figure 2.** (D) shows the overlay of three channels.

**Figure 5.**
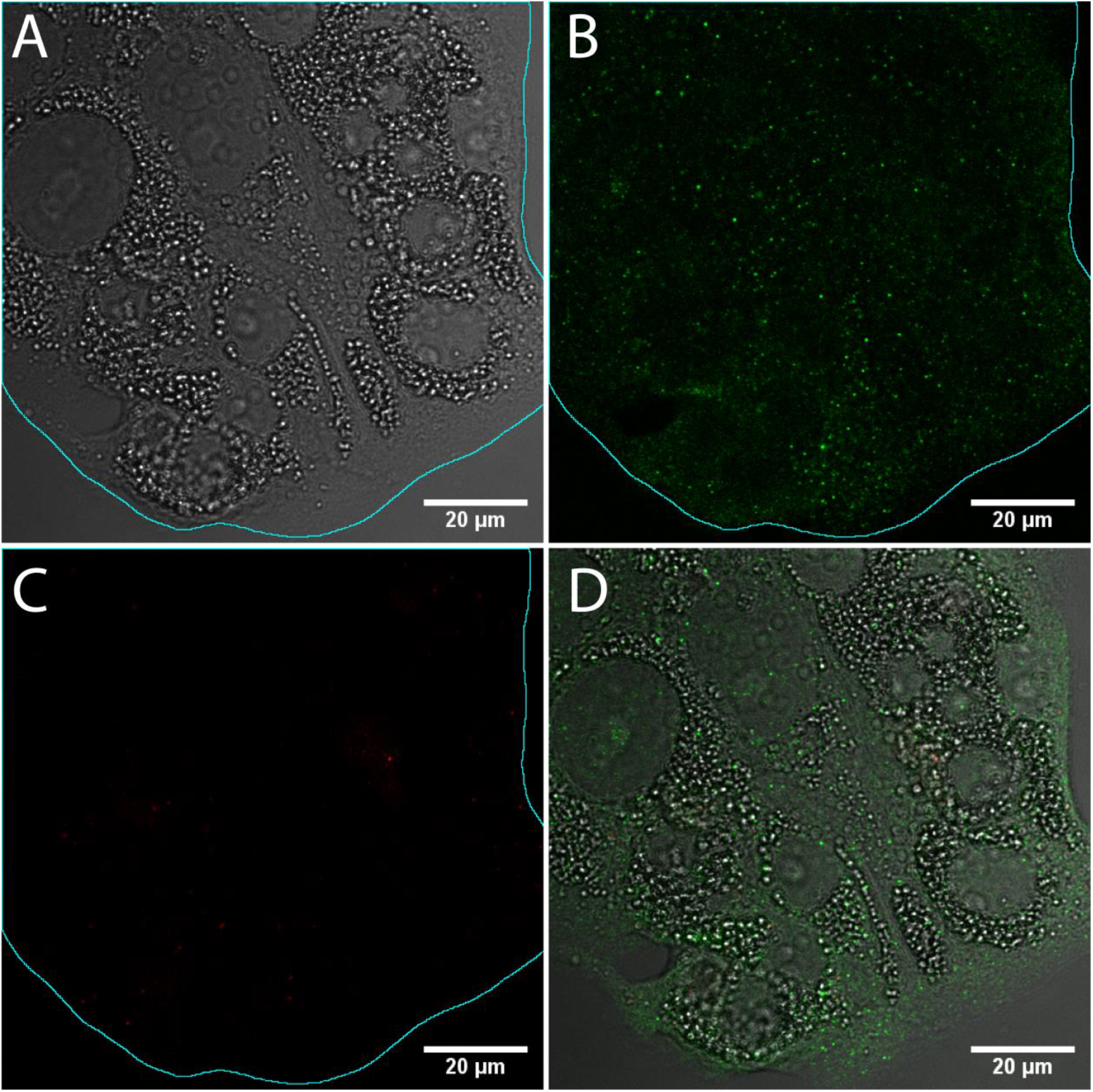
FND-loaded cells, cultured in butyrate-supplemented DMEM-HG, day 13. In the DIC image (A), a flat cluster of spread cells can be seen. The cell morphology is clearly different from the first day of incubation **(Figure 3)** or that of the non-treated cells **(Figure 4).** In the FITC channel (B), one can see increased fluorescence intensity of the cell cluster (cyan outline), indicating higher villin expression. The cells still carry FNDs in the cytoplasm (C). (D) shows the overlay of three channels.

**Figure 6.**
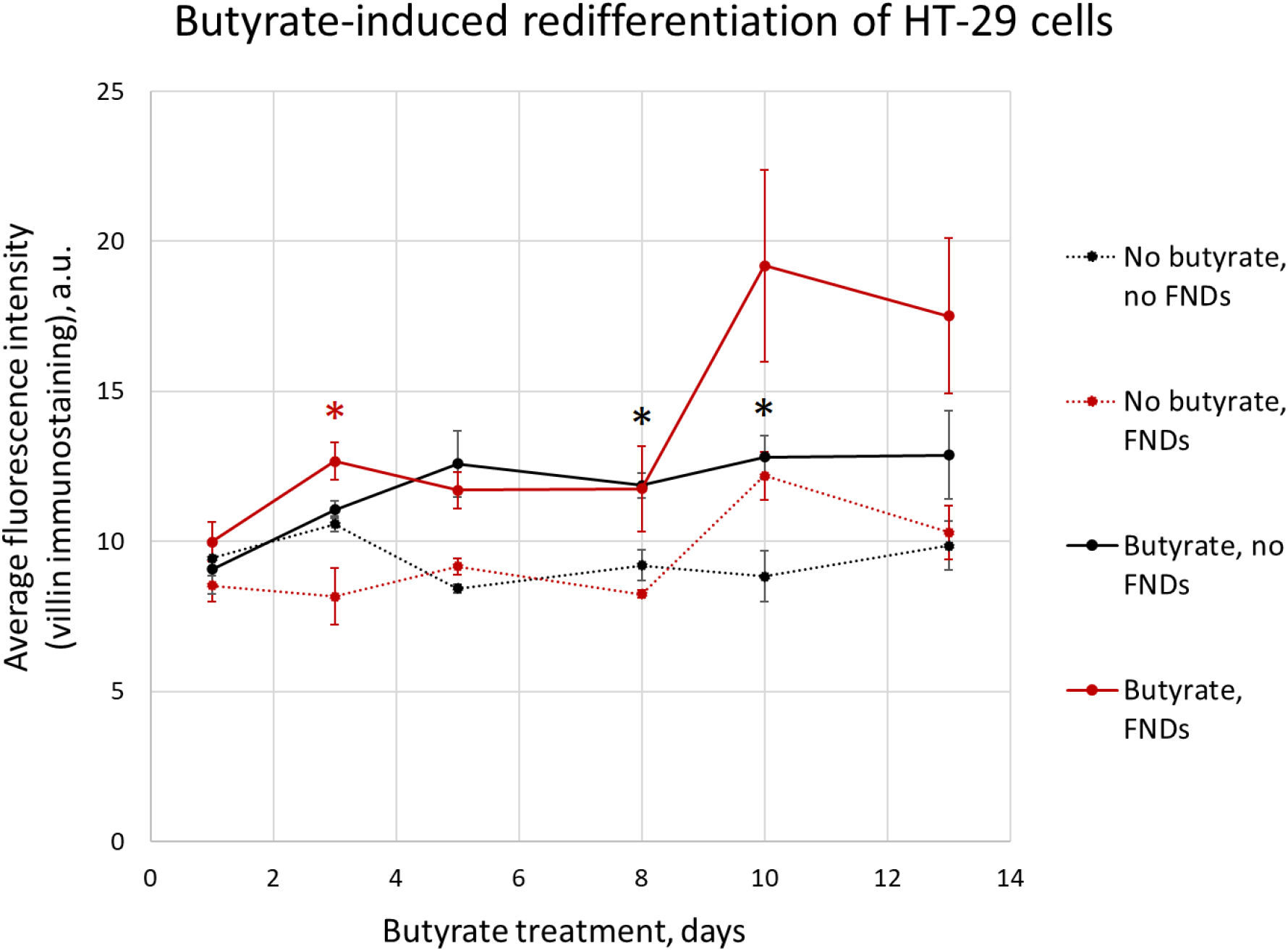
Villin expression in HT-29 cells, as determined by immunocytochemistry and image analysis. Each point is an average of at least 3 different fields of view within the sample. Error bars designate standard deviation. The statistical significance of the differences was tested with two-way ANOVE **(Table 1),** followed by Bonferroni’s multiple comparisons test. The multiple comparisons were performed separately within each experimental set (“no FNDs” – black asterisks; “FNDs” – red asterisk). The asterisks designate the differences between the groups with butyrate (solid lines in the plot) and without butyrate (dashed lines in the plot) at the same timepoint.

**Table 1.** Statistical analysis of villin expression in HT-29 cells. Two-way ANOVA was performed separately for cells with and without FNDs, to assess the impact of cell culture time and butyrate treatment.

| *Predictor variable* | **HT-29 cells without FNDs** |  | **HT-29 cells with FNDs** |  |
| --- | --- | --- | --- | --- |
|  | **p-value** | **% of total variation** | **p-value** | **% of total variation** |
| *Cell culture time (days)* | *<br>(0.0389) | 14.47 | ***<br>(0.0003) | 44.47 |
| *Sodium butyrate treatment* | ***<br>(0.0003) | 46.98 | ***<br>(0.0009) | 37.74 |
| *Interaction of time and butyrate* | ***<br>(0.0002) | 25.25 | *<br>(0.0117) | 7.888 |

Interestingly, FND-loaded HT-29 cells seemed to have higher villin expression than FND-free cells, regardless of butyrate treatment **(Figure 6).** While the differentiation process was more pronounced in butyrate-treated cells, time alone was a powerful predictor of villin expression **(Table 1).** These findings might indicate that the presence of FNDs in the cytoplasm affects the differentiation of HT-29 cells, even though it does not prevent this process from happening.

As can be clearly seen from the confocal images **(Figure 2–5)**, HT-29 cells retain FNDs in the cytoplasm even after 13 days in culture. This has allowed us to perform T1 measurements on the internalized nanodiamonds at different timepoints.

### 3.3. Internalized FNDs can be used to assess the free radical load in butyrate-treated and control cells

As HT-29 cells were cultured over the course of 13 days in butyrate-free medium, the average T1 values recorded from the cytoplasm would steadily go up **(Figure 7A).** The first statistically significant increase in T1 could be detected as early as at day 8 of culture. As the reported data were collected from different cells, FNDs, areas within the sample, and subcellular locations, these findings represent the change on the cell population level. Surprisingly, butyrate-treated cells showed stable T1 values over the course of the whole experiment **(Figure 7B).** These values were at the same level as those of non-treated cells at the earlier stages (**Figure 7C,** day 3-8), and significantly lower than in non-treated cells, starting from day 10. Thus, more differentiated HT-29 cells showed higher free radical load than undifferentiated cells, but only if the undifferentiated cells were cultured for a prolonged period of time. These findings are consistent with the electron spin resonance data from earlier studies, which showed that undifferentiated HT-29 cells have lower free radical load than butyrate-pre-treated cells^22^.

**Figure 7.**
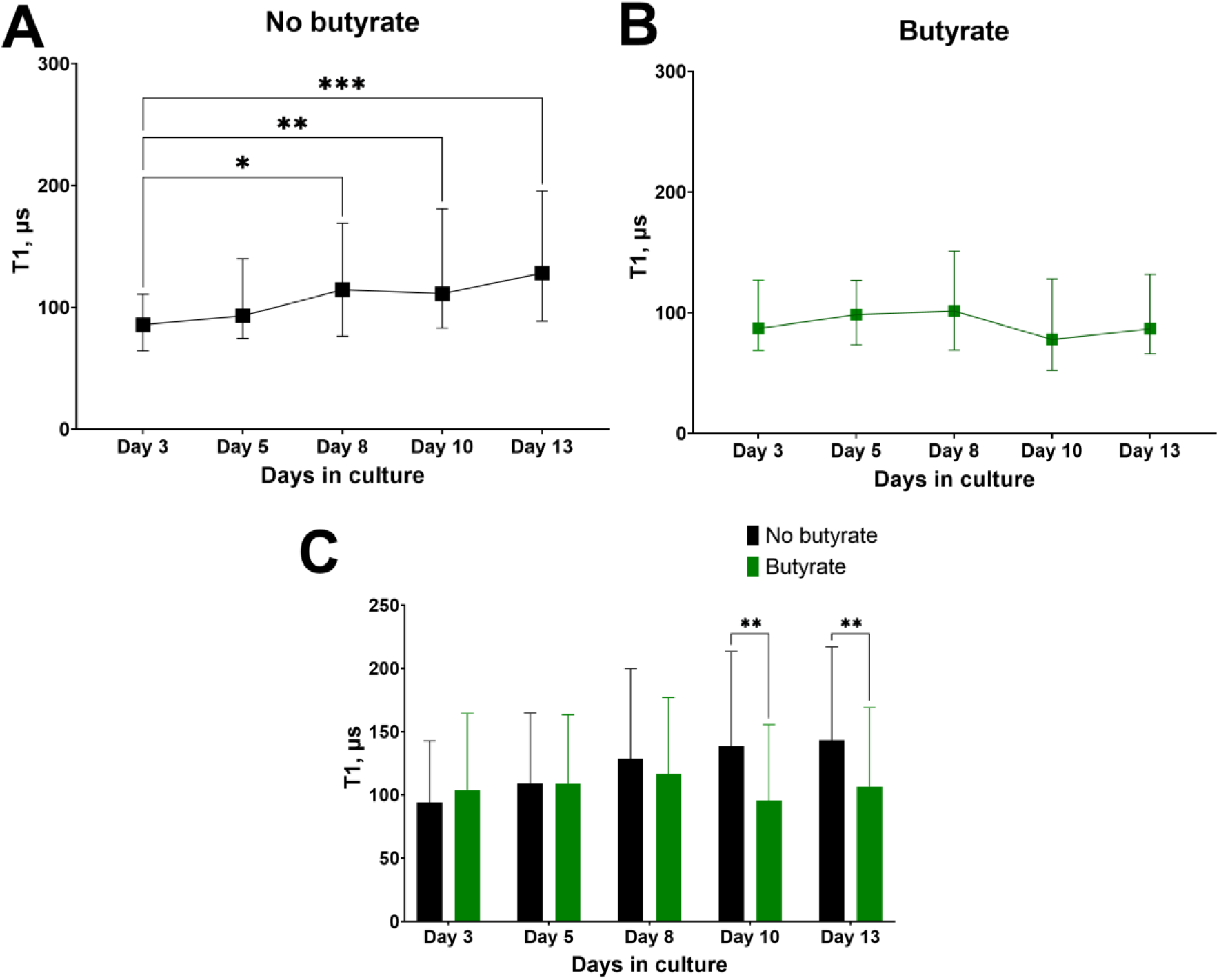
T1 relaxation of internalized FNDs in HT-29 cells, cultured in presence or absence of sodium butyrate. (A) HT-29 cells, cultured in butyrate-free medium, show progressively higher T1 values, indicating lower radical load. (B) If the cells are cultured in presence of sodium butyrate, T1 values remain the same throughout the whole period of the experiment. (C) Butyrate-free cells show significantly higher T1 values, as compared to butyrate-treated cells, starting from day 10 of the experiment.

## 4. Discussion

In this study, we have successfully performed enterocytic differentiation of HT-29 cells, carrying FNDs in the cytoplasm. Treating the cells with sodium butyrate results in the changes in cell morphology and levels of villin expression, which is consistent with previous findings^22^. Notably, we did not observe pronounced cell death in any of the samples at any of the experimental stages. At the same time, butyrate is known to cause cell cycle arrest and cell quiescence^20^, which can result in cells being less metabolically active. More extensive studies, including the assays to determine cell viability, cell redox state and intracellular production of reactive species, would thus be necessary to support the findings in this paper.

Interestingly, FND-loaded HT-29 cells seemed to have been pushed towards differentiation, regardless of the butyrate treatment. It has been previously demonstrated that the presence of FNDs in the cytoplasm does not impede the differentiation of various cell types, including the differentiation of pre-adipocytes and mesenchymal stromal cells into adipocytes^10,11^, adipose-derived stem/stromal cells into chondrocytes^12^, embryonal cancer stem cells into neurons^13^, and lung progenitor cells into pneumocytes^14^. To our knowledge, this is the first study demonstrating the possibility of enterocytic differentiation of the cells, carrying FNDs. However, FNDs can be more than just inert particles: for example, neural stem cells grown on the substrate, coated with nanodiamond films, spontaneously differentiate into neurons and exhibit higher neurite density^27,28^. Similarly, the osteogenic differentiation of osteoblast-like cells grown on nanocrystalline diamond films is more efficient, as compared to polystyrene substrate^29^. At the same time, little is known about the beneficial effects of internalized FNDs on the cell differentiation. More research is thus needed to test, whether FNDs indeed promote this process and what mechanisms might be involved.

Consistently with previous studies, butyrate-treated cells showed distinctly higher levels of free radical load, as compared to the non-treated cells, but only after a prolonged incubation with butyrate. Interestingly, we observed changes in the control sample, and not in the butyrate-treated sample. Our data suggest that cultured HT-29 cells need approximately 8 days to develop the “higher T1” status, and that freshly seeded cells show low T1 values, comparable with those of butyrate-treated cells. While the results of our measurements are not directly affected by the number of cells in the sample (unlike in case of fluorescent/colorimetric plate-based assays, such as DCFDA or MTT), higher cell counts can affect the physiology of individual cells. Among other things, the oxygen content in the sample would be lower, if the cell counts are high, which could create the hypoxic zones. On the one hand, reduced oxygen content will result in reduced amount of reactive oxygen species formed in the cell. On the other hand, it is well known that, in cancer cells, hypoxia leads to oxidative stress and switch to glycolysis, but this, in turn, allows the cell to ramp up the antioxidant production and to mitigate the oxidative damage^30^. Due to the combination of these factors, cancer cells in the hypoxic conditions might exhibit lower free radical load than either differentiated cells or cells in a normoxic environment. To account for the effects of the cell proliferation, resulting in a crowded environment, future experiments would be needed, in which the cell density would be matched between butyrate-treated and non-treated samples.

The observed discrepancy in free radical load can also be an inherent quality of the cells, which depends on their differentiation status and not on the environmental factors. If these results are supported by additional experiments, our findings can pave the way for using nanodiamond relaxometry to distinguish between more and less differentiated cells. This can be useful to get a deeper insight into the single-cell physiology of differentiating cells and can have powerful implications for stem and cancer cell biology. At the same time, diamond relaxometry could be transformed into a diagnostic tool, allowing to estimate the differentiation status of the tumour cells in clinical samples, which is closely linked to the malignancy of the tumor^31^. As our method, in principle, allows to track the free radical levels in individual cells, it can also be used to assess the responsiveness of the patient-derived cells to the proposed treatment. Knowing the differentiation status of the cells in the sample, one can specifically look at the less differentiated (and more malignant) cells to test the therapy resistance. This approach can thus contribute to the development of more precise patient-tailored approaches to cancer therapy.

## 5. Conclusions

This study reports on the successful differentiation of HT-29 cells, pre-loaded with FNDs. Over the course of the 13-day treatment with sodium butyrate, the cultured cells developed morphological and molecular traits, characteristic of normal enterocytic differentiation. This process was not impeded by the presence of nanodiamonds in the cytoplasm; in fact, non-functionalized FNDs alone showed signs of promoting enterocytic differentiation. The particles were successfully retained in the cytoplasm throughout the experiment. FNDs could be identified inside the HT-29 cells at all timepoints, which allowed us to use them for quantum-based measurements of the intracellular free radical load.

Non-differentiated cells showed an increase in T1 values (corresponding to decrease in free radical levels) over the course of the experiment. Butyrate-treated cells, on the contrary, maintained relatively low T1 values (high free radical load) throughout the differentiation process. By day 10 of the experiment, butyrate-treated cells showed significantly lower T1 values than non-differentiated, non-treated HT-29 cells. These findings are consistent with the previously reported results on the free radical load in these two subpopulations^22^. However, our data suggest that it is the free radical load in non-differentiated cells that is altered during the cell culture. Further research would be needed to establish whether these changes are inherent to the cells and reflect their differentiation status or are caused by the environmental factors, such as the oxygen levels and nutrient availability. If free radical levels indeed depend on the differentiation stage of the cell, our approach can be used to better distinguish between more and less differentiated cells within the same sample, both in the fundamental research and the clinical setting.

